# Machine learning meets classical computer vision for accurate cell identification

**DOI:** 10.1101/2022.02.27.482183

**Authors:** Elham Karimi, Morteza Rezanejad, Benoit Fiset, Lucas Perus, Sheri A. C. McDowell, Azadeh Arabzadeh, Gaspard Beugnot, Peter Siegel, Marie-Christine Guiot, Daniela F. Quail, Kaleem Siddiqi, Logan A. Walsh

**Affiliations:** Rosalind and Morris Goodman Cancer Research Centre, McGill University, Montreal, QC, Canada; School of Computer Science & Centre for Intelligent Machines, McGill University, Montreal, QC, Canada; Departments of Psychology and Computer Science, University of Toronto, Toronto, ON, Canada; Department of Physiology, Faculty of Medicine, McGill University, Montreal, QC, Canada; Inria, École Normale Supérieure, CNRS, PSL Research University, Paris, France; Department of Medicine, Division of Experimental Medicine, McGill University, Montreal, QC, Canada; Department of Biochemistry, McGill University, Montreal, QC, Canada; Department of Pathology, McGill University, Montreal, QC, Canada; Department of Human Genetics, McGill University, Montreal, QC, Canada

## Abstract

High-parameter multiplex immunostaining techniques have revolutionized our ability to image healthy and diseased tissues with unprecedented depth; however, accurate cell identification and segmentation remain significant downstream challenges. Identifying individual cells with high precision is a requisite to reliably and reproducibly interpret acquired data. Here we introduce CIRCLE, a cell identification pipeline that combines classical and modern machine learning-based computer vision algorithms to address the shortcomings of current cell segmentation tools for 2D images. CIRCLE is a fully automated hybrid cell detection model, eliminating subjective investigator bias and enabling high-throughput image analysis. CIRCLE accurately distinguishes cells across diverse tissues microenvironments, resolves low-resolution structures, and can be applied to any 2D image that contains nuclei. Importantly, we quantitatively demonstrate that CIRCLE outperforms current state-of-the-art image segmentation tools using multiple accuracy measures. As high-throughput multiplex imaging grows closer toward standard practice for histology, integration of CIRCLE into analysis protocols will deliver unparalleled segmentation quality.

---

Recent advances in multiplex imaging have revolutionized our ability to identify functionally meaningful cell populations among spatially and phenotypically heterogeneous tissues. Single-cell profiling using imaging mass cytometry (IMC), co-detection by indexing (CODEX), DNA-barcoding, serial immunostaining, or similar high-parameter imaging approaches have revealed that clonal evolution and spatially distinct tissue microenvironments drive cellular heterogeneity and control pathophysiology [1, 2, 3, 4, 5, 6, 7]. For example, the recent systematic, multi-dimensional interrogation of breast cancer using IMC revealed a detailed spatial map of single-cell phenotypes and cellular communities and resolved novel subtypes of breast cancer with distinct clinical outcomes [2]. Such multicellular spatial information provides a basis to study how geographical and phenotypic tissue features influence disease outcomes.

The most important requirement of image-based cellular profiling is the ability to identify individual cells through segmentation with extremely high precision, as this directly dictates the correctness and clinical relevance of all downstream discovery-based analyses. To do this, nuclei must be accurately distinguished, therefore the value of the technology hinges on accurate nuclear segmentation methodologies. This is a fundamental challenge in biomedical imaging primarily due to: (i) the inherently low contrast of edges and geometric boundaries, (ii) the lack of color in the image (most imaging technologies output in grayscale and use pseudo color), (iii) the lack of a unique defining texture, (iv) the varying shapes and sizes of cells, (v) the presence of overlapping cells in the image due to partial volume effects, (vi) the lack of ground truth examples, which limits the performance of machine learning methods (due to the inherent lack of training data), and (vii) the high density of cells, particularly in diseased tissue such as tumors. Each of these properties affects the design of computer vision algorithms to segment cell instances. It is also largely impossible to manually verify image quality in high-throughput experiments and there is a lack of automated methods to objectively flag or remove images and cells that are affected by artifacts. As a result, the existing tools for image-based cellular profiling regrettably require significant subjective manual intervention, which limits data reproducibility and reliability.

Over the years several methods have been developed for image-based cell identification, but each approach has limitations that are often overlooked. Image thresholding while adapting to local contrast [8] is very common. Here regions are distinguished by intensity averages or other measures of separation that are chosen either heuristically or intelligently. While most cell profiling techniques can benefit from this as an initial step, there is no guarantee that a distinct separation of cells can be achieved. Another classical method for medical image segmentation is the watershed algorithm, which equates pixel values to local topography [9, 10, 11]. Watershed based algorithms are widely used and are very effective when the regions to be segmented have homogeneous intensities. However, they often fail to differentiate between texture-based edges and true organelle boundaries. Active contour models [12, 13] allow geometric curves or surfaces to evolve and capture the boundaries of specific visual elements. They do so by minimizing a suitable energy functional, which incorporates both local (boundary) and global (region) image information so that the curves or surfaces converge to object boundaries [14, 15]. However, the outputs obtained by such models can be sensitive to the choice of initial seed contours, and separating overlapping cells can be a challenge. A further obstacle in using active contour models is the challenge of finding initial seed mask contours.

With advances in machine learning-based models, several promising approaches to image-based cell identification have been developed in both semi-supervised and supervised settings [16]. Many of these models use existing popular deep learning frameworks such as UNet [17] and MaskRCNN [18]. A general requirement of deep neural network (DNN)-based models is the availability of large annotated training datasets. Further, DNN models represent somewhat of a ‘black box’ for the user since they presently offer little insight for troubleshooting when a desired cell population is not accurately identified. Despite these limitations, DNN models can benefit from a concept referred to as “transfer learning” [19, 20, 21, 22]. When deep models are initially trained on huge datasets of generic objects they learn to extract certain high-level visual features that aid a recognition or categorization task. Their pre-trained weights can then be later fine-tuned on specific tasks, with smaller amounts of task-specific training data.

In recent years, additional supervised learning-based cell identification methods from data points that represent cell locations have been established. There are advantages to approaching cell identification by combining both region and boundary information. Advances in UNet-based models enable advanced cell identification mechanisms in which all pixels that are surrounded by a cell instance are marked and labeled separately. Among these methods, StarDist[23] is an effective UNet-based model that learns a shape representation for cells when it detects them in images. The UNet backbone contributes to excellent foreground detection[24] and as a result, StarDist has a high recall, successfully finding locations where candidate cells exist. On the other hand, the cell boundaries provided by StarDist are not always accurately placed. In particular, its precision suffers when the underlying images have a low signal-to-noise ratio. Another recent method which also uses deep neural networks, Cellpose [25], promotes the benefits of crowdsourcing to acquire labeled training datasets to gradually improve the performance of such methods. Motivated by the strengths and weaknesses of the aforementioned strategies for image-based cell-profiling, we developed CIRCLE (Cell segmentatIon acRoss sCaLEs), a state-of-the-art framework that is capable of automated processing of high dimensional images with exceptional accuracy (Fig. 1). CIRCLE is a 2D cell identification pipeline that combines classical and modern machine learning-based computer vision algorithms to address shortcomings of commonly used image segmentation tools ([25, 18, 23, 10, 26, 9]). CIRCLE integrates a CNN for nuclear segmentation, with a novel combined active contour model and scale selection process. The CIRCLE pipeline can accommodate low-resolution images and is implemented efficiently.

**Figure 1:**
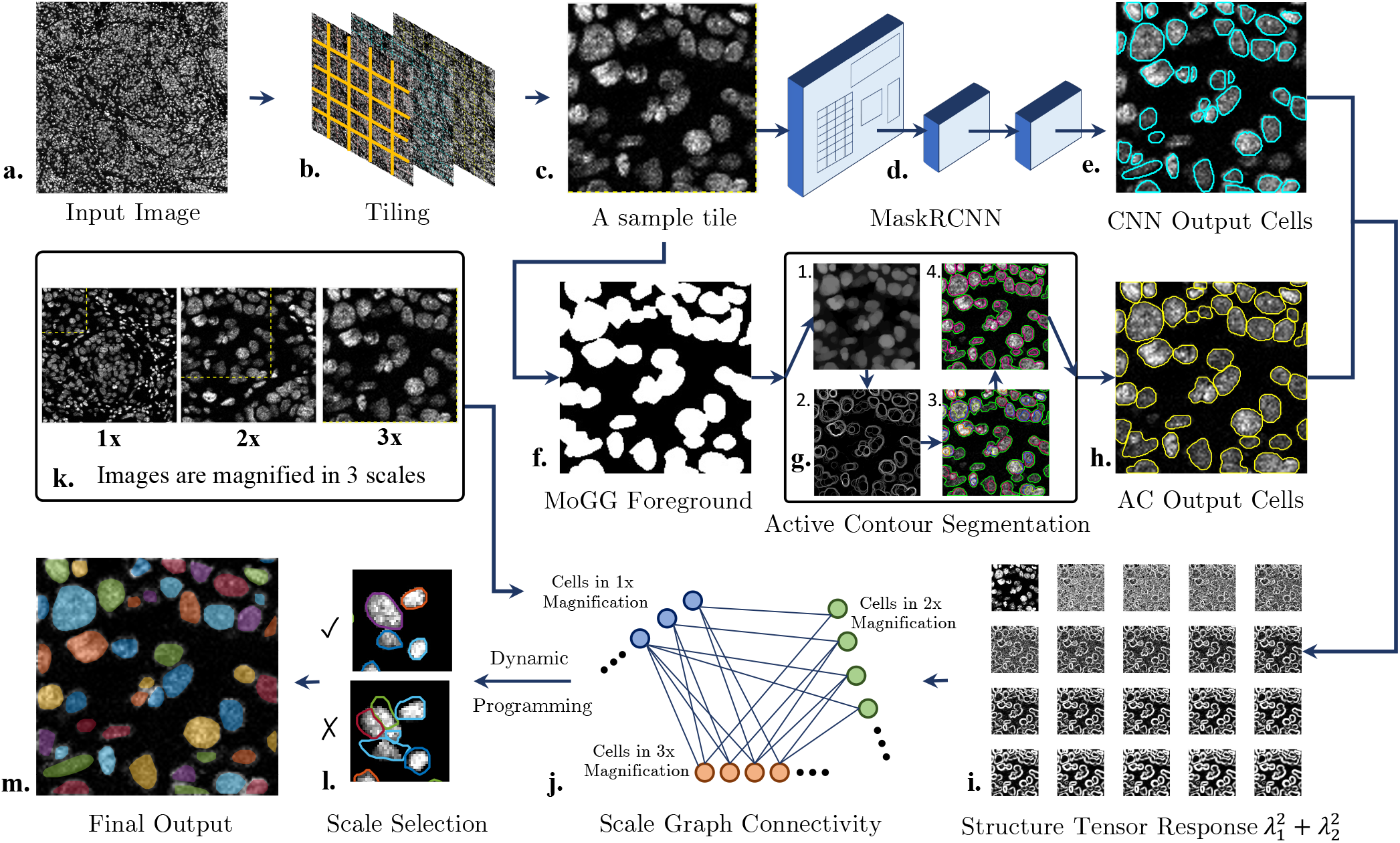
Schematic overview of the CIRCLE pipeline. **a**. Representative input image (stained nuclei). **b**. Image is tiled, and shifted copies of the tiles are created so that cells lying at the borders are not missed. **c**. Each tile is processed at several magnification scales (represented in **k**.) allowing cells of varying sizes to be detected. **d-e**. For the deep model, MaskRCNN is used for instance cell segmentation. **f**. Concomitantly, tiles also serve as inputs for Mixture of Generalized Gaussian (MoGG) foreground mask generation. **g**. Active contour based refinement is applied to the MoGG foreground mask to yield a cell segmentation result for each tile that is independent of MaskRCNN **h**. The MaskRCNN and active contour output cells are pooled together. **i**. For each cell a structure tensor based feature is computed within a narrow band of each cell boundary. **j**. A scale graph is built to consider the structure tensor based feature across different magnification scales. **l**. Using dynamic programming, we select cells amongst the different magnifications. For overlapping detections, the best magnification is chosen and the other detections in the pooled results are removed. **m**. Representative image of the final output.

We first divide the input image into smaller tiles that can each be analyzed separately. We detect cells in each of these smaller tiles at multiple magnifications, using both a deep neural network model and an active contour segmentation method. For the former process, we use the state-of-the-art Mask Region Convolutional Neural Network (MaskRCNN) architecture [18] to generate segmentation masks for each cell.

MaskRCNN provides high precision and requires minimal complex post-processing. The same image patch is also provided as input to a novel active contour segmentation method (ACSM). In the ACSM, the foreground regions are first found using a Mixture of Generalised Gaussians algorithm to include nuclei that could be potentially missed by the DNN model. We then apply a series of image processing operations to find all possible cells within these foreground regions. At the end of this process, we create a set of structure tensor (ST) maps to describe the intensity distribution in the vicinity of the estimated cell boundaries at each magnification scale. We then construct a connectivity graph across the detected cells at each scale. Using features derived from the ST, we apply a dynamic programming method to select the best scale for each cell, like the one that maximizes the ST feature response at the boundary. Scale selection enables our algorithm to avoid under-segmentation and over-segmentation of cells, even in very noisy conditions, and to handle cells of a variety of sizes.

To quantitatively compare the accuracy of CIRCLE against current state-of-the-art segmentation methods, first, we trained and fine-tuned the deep models CIRCLE, MaskRCNN, and StarDist on the 2018 Data Science Bowl (DSB 2018) competition dataset (https://www.kaggle.com/c/data-science-bowl-2018). Cellpose was pre-trained and fine tuned on multiple large datasets including the DSB 2018. Next, we created an entirely new test dataset, GCIMC, by meticulously manually segmenting four 1000 × 1000 pixel images generated through IMC of brain metastases from 4 patients. GCIMC contains **12692** segmented nuclei. When applied to the GCIMC dataset, CIRCLE was the most accurate segmentation method as determined by several measures, including 1) average precision across a range of intersection-over-union (IoU) thresholds (Fig. 2, a.), 2) precision versus recall (Fig. 2, b.) and 3) the Dice Coefficient (Fig. 2, d.). To qualitatively visualize the accuracy, dynamic range, and broad applicability of CIRCLE, we have included representative segmented IMC images from normal and cancerous tissues that exhibit various challenging features including poor contrast, varying cell size, and overlapping cells. (Fig. 3). These data highlight that although current methods may perform reasonably well on select tissues, CIRCLE uniformly is the most accurate, especially when confronted with heterogeneous and complex tissue microenvironments.

**Figure 2:**
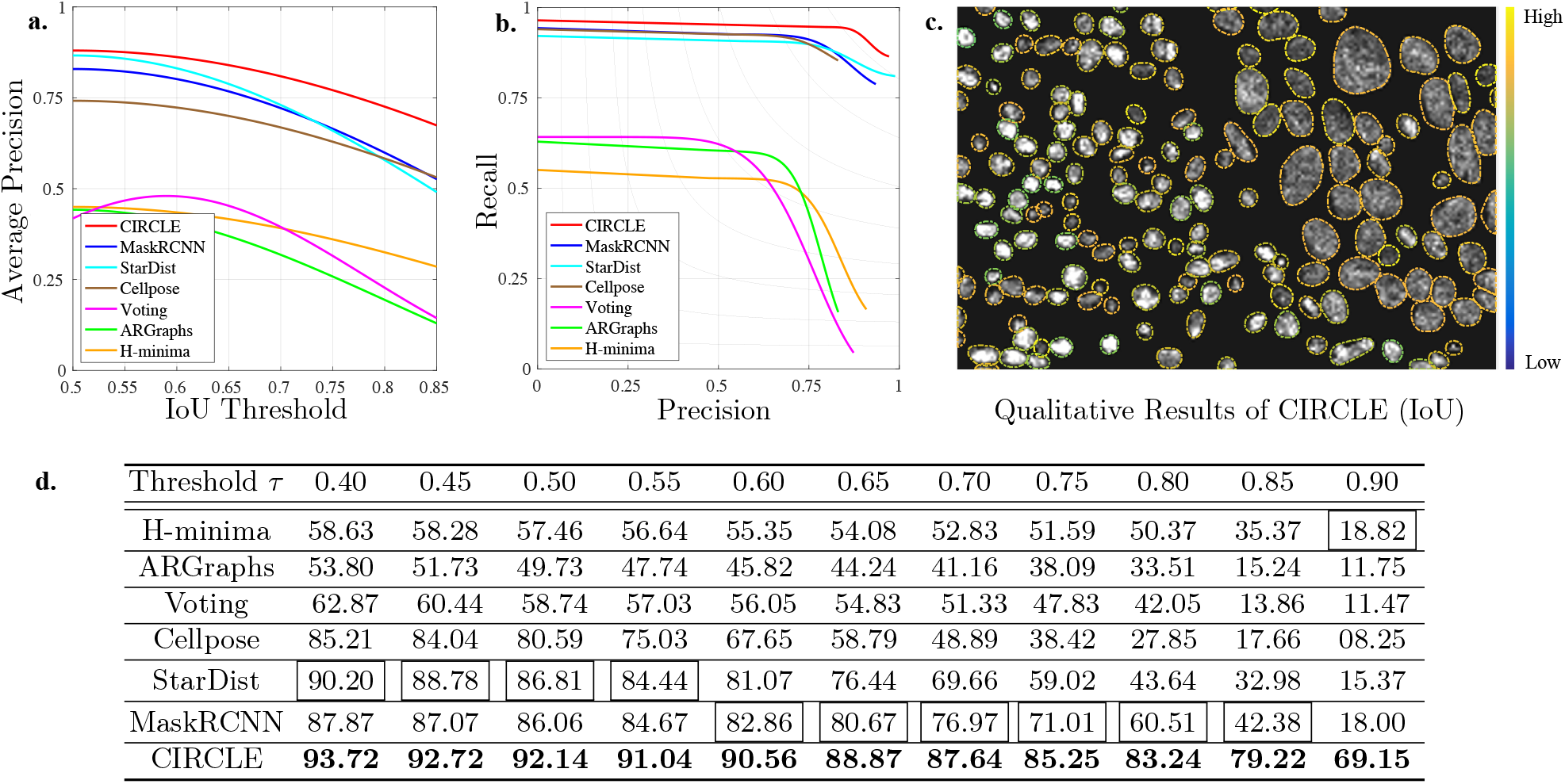
Quantitative comparison of CIRCLE with current state-of-the-art methods. We have applied CIRCLE to low-resolution multiplexed IMC data (GCIMC dataset) and have compared its accuracy to common segmentation algorithms. **a**. Average precision 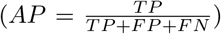. **b**. Precision versus Recall. **c**. Representative output of CIRCLE segmentation (boundaries are colored from blue to yellow according to their *several intersection over union* (IoU) values ranging from 0.5 to 1.0). **d**. Quantitative assessment of segmentation accuracy (Dice Coefficient, defined using precision (P) and recall (R) as 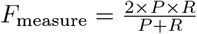 for IoU thresholds *τ*).

**Figure 3:**
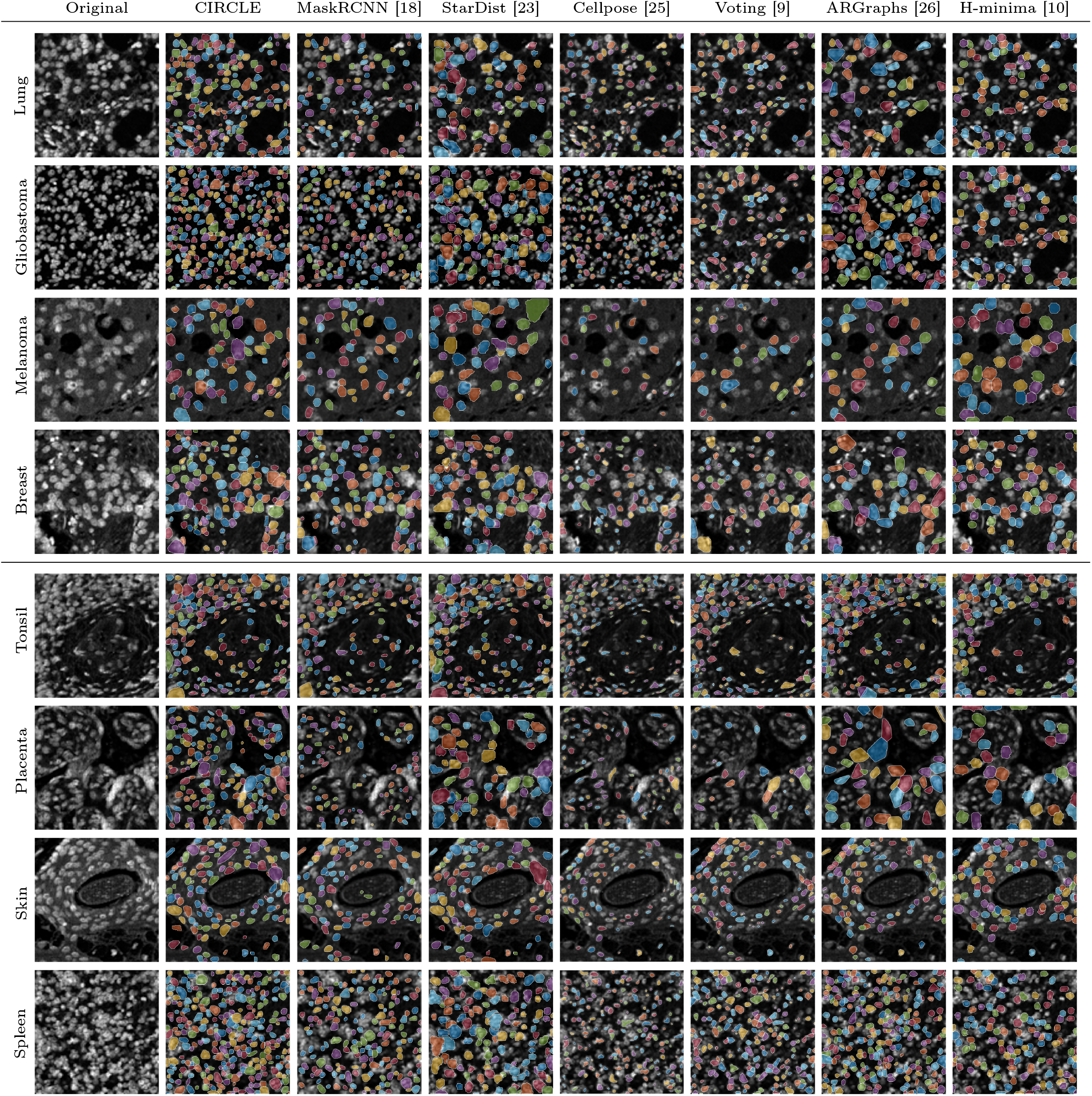
Qualitative comparison of CIRCLE with current state-of-the-art methods. Representative IMC images from normal and cancer tissues, reflecting a range of challenging cases including poor contrast, varying cell size, and overlapping cells, segmented with the indicated methods.

We are currently in an imaging revolution where images have become an unbiased source of quantitative information for biological phenotypes. As more studies exploit new technologies capable of generating highly multiplexed landscapes of tissue with spatial resolution, the need for accurate segmentation will only grow. In this setting, CIRCLE provides precise and automated cell identification while overcoming several limitations of other tools, offering a fully automated hybrid cell detection model to segment nuclei in 2D images.

## Methods

We describe a number of components of our workflow in the subsections below. These methods are combined in the manner illustrated in Figure 1. The general idea is to consider both the output of a powerful CNN-based segmentation method (Section 2) and one that uses a foreground detection approach first (Section 3), followed by an active contour refinement stage to separate potentially overlapping cells (Section 4). The two independent sets of candidate cell segmentations are then pooled and among these a selection criterion is applied across different magnification scales to select the best scale for those detections that have a spatial overlap (Section 5). This scale selection process allows for variation in cell size.

### 1. Tiling

To make our approach memory efficient, we divide each input image into a smaller set of tiles. The tiles are chosen to be larger than the maximum possible size of a cell to be detected. Tiling facilitates the use of standard CNN implementations, which typically assume that the input images are small, and mitigates the effect of non-uniform luminance across larger windows. Assuming that the source image is of size *W* × *H* and that the tile width is *w*_*s*_, we partition the image into a grid with *m* × *n* tiles, where: 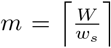 and 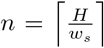. The tile width parameter is chosen empirically, e.g., for the images that are 1000 × 1000 in size, we chose *w*_*s*_ to be 200 to yield 200 tiles. Each tile is provided as a separate input to our method. All the detected cells that intersect a tile boundary are discarded. To ensure that no cells are missed we create two copies of the original tiling, shifting it by 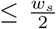 to the South and to the East, and use these shifted tiles as inputs as well. The choice of *w*_*s*_ should be no smaller than the maximum possible size of a cell. To avoid duplicate detections of the same cell due to the additional shifted tiles, we maintain an occupancy image, which is the size of the original image, and fill it with the area that each detected cell covers.

### 2. MaskRCNN Cell Detection

To obtain one set of candidate cells, we apply the Mask Region Convolutional Neural Network (MaskRCNN) architecture to segment cell nuclei [18] (Fig. 1d-e). This network provides advantages over the popular UNet approach [17], having evolved through four different earlier versions. The latest version uses an RoIAlign layer to correct for misalignments and generates both the bounding boxes and segmentation masks for each cell in the image. MaskRCNN is already used [27, 28, 29, 30, 31, 32] to segment nuclei/cells or lesions in medical imaging. We used the existing model, pre-trained on the COCO dataset [33], as a starting point and then fine-tuned it on the data science bowl public dataset available at https://www.kaggle.com/c/data-science-bowl-2018. We applied the various standard data augmentation techniques including both proportional and non-proportional image resizing, random rotations in increments of 10 degrees (36 rotations in all), and vertical and horizontal flips. The fine-tuning consists of training the network in 3 batches of 25 epochs each, where each batch’s trained weights were used as initial weights for the next batch, and the learning rates were reduced for the subsequent batch.

### 3. Foreground Detection

Foreground detection refers to the marking of any region in the image that could potentially contain a cell. Whereas a naive approach would involve intensity-based thresholding, more sophisticated methods use histograms [38, 39, 40, 41, 42], clustering [43, 44, 39, 45], or the entropy of foreground and background regions [46, 47, 48, 49, 34]. Since our main purpose is to mark regions whose boundaries will be later refined, a multiclass histogram thresholding approach to efficiently model multi-modal class-conditional distributions using mixtures of generalized Gaussian distributions (MoGG) [50] works well, in comparison to alternative approaches including an entropy-based method [34], the use of an adaptive threshold [35], and the use of superpixels for oversegmentation [36, 37] (Extended data Figure 1). MoGG provides a flexible and suitable tool for various problems in image segmentation including ones that arise in medical image analysis [51, 52, 53, 54, 55]. The foreground detection is an essential step since it improves recall by adding additional regions that may have been missed by the neural network. These foreground regions are refined using an active contour model to provide candidate cell segmentations, as described next.

### 4. Active Contour Based Cell Segmentation

The output of the MoGG foreground detection method is now refined to better separate potentially overlapping cells. We exploit the property that when two or more convex objects overlap in the 2D plane, there are two local peaks in the distance function from their combined boundary.

#### 4.1. Morphological Opening

Morphological opening consists of an erosion step followed by a dilation step [56] to remove structures smaller than the diameter of the structuring element: *I*_*o*_ = *I* ○ *s* = (*I* ⊖ *s*) ⊕ *s*. Here *I* is the input image patch, and *s* is the structuring element. We use a circular structuring element with a diameter *d* chosen to be linearly proportional to the considered scale. The result of this opening operation on the sample tile is shown in Figure 1g, panel 1. The opened image results in interior cell regions with a more homogeneous appearance.

#### 4.2. Gradient Rings

We now devise a way to separate overlapping cells in the opened image. We use the magnitude of the gradient within the opened image as a measure of cell boundary strength (Fig. 1g, panel 2): 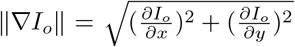 where 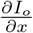 and 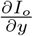 are the partial derivatives of the image in the *x* and *y* directions, respectively. We then consider level sets of the gradient magnitude, normalized to lie between 0 and 1, to provide a series of nested gradient rings, as illustrated in Figure 1g, panel 3.

#### 4.3. Generating Seed Contours

We now process the nested gradient rings in order to provide seed contours to segment individual cells. Each gradient ring, corresponding to a simple closed contour, is treated separately (as illustrated using different colors in Fig. 1g, panel 3). We denote the *i*^*th*^ ring by *r*_*i*_ and prune rings in a recursive manner. Here, ring *r*_*i*_ is removed if there is another ring *r*_*j*_ such that *r*_*i*_ ⊆ *r*_*j*_ and no other ring *r*_*k*_ exists such that *r*_*k*_ ⊆ *r*_*j*_. The pruning stops when no further rings can be removed. We verify the *r*_*i*_ ⊆ *r*_*j*_ condition by checking whether the area bounded by *r*_*i*_ is contained within the area bounded ring *r*_*j*_. The surviving rings are shown in red in Figure 1g, panel 4.

#### 4.4. Active Contour Cell Segmentation

In a final step, we refine the surviving rings to segment likely cell boundaries within an image tile, using the Chan-Vese region-based active contour [57]. This formulation best separates each MoGG foreground region into two classes, under the assumption that each class is homogeneous in intensity and thus well represented by its mean. Here, we assume a system of multiple active contours that evolve simultaneously at each time step. Let *r* represent a ring, and *u*_0_ represent the set of pixels within the same MoGG foreground region, excluding *r*. The algorithm evolves the contour to minimize the following energy function, ultimately placing the evolved contour at the boundary of two classes:

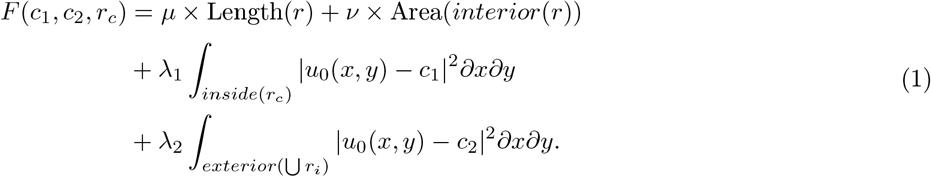

Here *interior*(*r*) refers to the pixels contained within *r*, and *exterior*(*r*_*i*_) is the set of all pixels in the foreground region excluding those associated with all present rings. *c*_1_ is the mean intensity of *u*_0_ inside *r* and *c*_2_ is the mean intensity of *u*_0_ in the region outside *r*, within the foreground region. *µ* ≥ 0, *ν* ≥ 0, and *λ*_1_, *λ*_2_ *>* 0 are empirically chosen fixed weight parameters. Using a level set formulation [58, 59, 60], we evolve the ring *r* so as to minimize the above energy. Each ring is updated using the following level set equation at each time step:

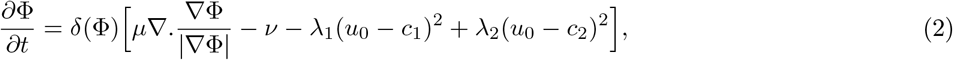

where *δ* is the Dirac-delta function. Here, Φ (*x, y, r*) is the embedding function whose level sets represent the evolving rings [58, 60, 59]. With multiple level sets evolving simultaneously, the pixels that are inside and outside of rings within *u*_0_ are updated in time. For the level set update we use the numerical method in [61]. The resulting detected cells are shown in Figure 1h.

### 5. Scale Selection

We now have two independent sets of candidate cells at each magnification scale: those obtained by the CNN (Fig. 1e) and those obtained by the active contour detection process (Fig. 1h). We have devised a method to select between these candidate detections over the considered scales. We first compute a salience score for each cell at each magnification scale. To do so, for each magnification, we blur the image patch with Gaussians over a range of blurring scales. We then compute the Structure Tensor (ST) for all standard deviations at each pixel. The ST is computed at each pixel **p** as:

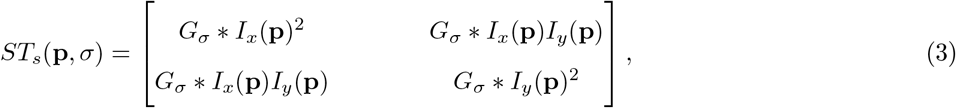

where *I*_*x*_(**p**) and *I*_*y*_(**p**) are partial derivatives in space in the directions *x* and *y* of the image *I*, computed using central differences. *G*_*σ*_ is a Gaussian filter of length 2*σ* + 1 with a standard deviation of 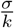, where *k* is an arbitrary positive constant.

We then compute a local feature descriptor measure using the sum of the squared eigenvalues of the ST matrix: 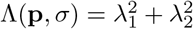. In principle, we expect 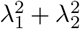 to be relatively large at a location near the cell boundary or when when two cell boundaries are touching each other (Fig. 1 (i)). For each scale *s* let *n*_*s*_ be the number of pooled CNN and active contour cell detections, stored in a list 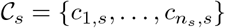. We define a salience score for each cell as the highest average value of Λ(**p**) within the vicinity of a boundary of a cell, amongst all possible standard deviations as

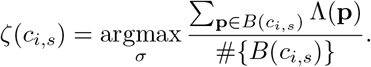

Here *B*(*c*_*i,s*_) is a one-pixel expansion of the cell boundary *c*_*s,i*_ towards the inside and outside. We accommodate for size variation in cell nuclei by considering 3 different magnification scales (Fig. 1k). To select between these magnification scales and also the different cell generation methods, we construct a connectivity graph across cells and magnification scales. Each cell at each magnification scale is a node of the connectivity graph, with a weight of *c*_*i,s*_. We connect two cells 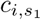 and 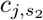 if the ratio of the common area between those two cells over the union of the areas of these cells is at above a certain threshold:

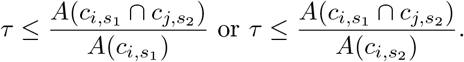

Here *A*(*c*_*k,m*_) represents the area of the *k*^*th*^ cell at magnification *m* and *τ* is a particular threshold chosen between 0 to 1. In our experiments, we chose *τ* to be 0.85. After the entire connectivity graph is constructed (Fig. 1j), we update each node’s *ζ*(*c*_*s,i*_) score by dividing it by the degree of that node: 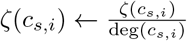.

Following the construction of the connectivity graph, we select cells using a dynamic programming algorithm that maximizes the sum of *ζ*(*c*_*s,i*_) scores over all identified cells. Let 𝒞_*i*_ represent the cells at *i*^*th*^ magnification level. Our goal is to optimize an energy function 𝒰(𝒞_1_, 𝒞_2_, 𝒞_3_) that selects cells between these magnification scales and the two detection methods, while avoiding overlapping detections. For this we construct the following energy

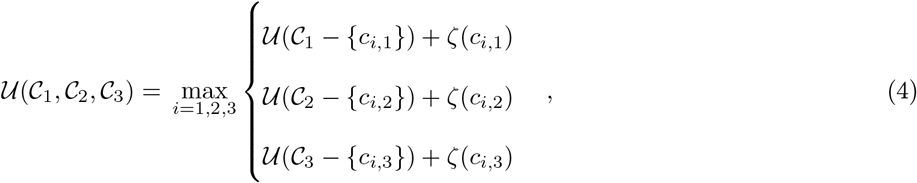

where 𝒞_*j*_ − {*c*_*i,j*_} refers to the removal of cell *c*_*i,j*_ and its connections from all cell sets 𝒞_*i*_. As the connections between overlapping cells from the two different methods and the different magnifications scales are sparse, we used the Cuthill-McKee algorithm [62] to permute the sparse matrix of connections, for increased computational efficiency.

## Data and Code Availability

All data and source code will be made available without restrictions. Ethics approval IRB00010120. Normal tissues were de-identified excess diagnostic tissue controls that were slated for incineration.

---

**Extended data Figure 1.**
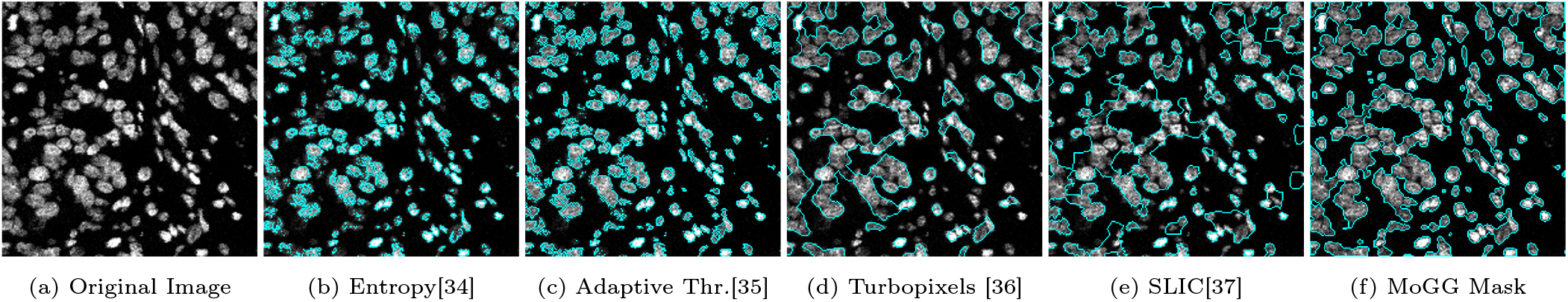
A comparison of foreground detection approaches on a typical tissue sample. In general we found the MoGG method to be robust to non-uniformities in luminance and degradation in image quality, and to have a high ratio of true positives to false negatives (missed foreground detections).

